# Identification of an *ERCC2* mutation associated mutational signature of nucleotide excision repair deficiency in targeted panel sequencing data

**DOI:** 10.64898/2026.02.17.706196

**Authors:** Olivera Stojkova, Judit Börcsök, Zsofia Sztupinszki, Miklos Diossy, Aurel Prosz, Alexander Neil, Kent W. Mouw, Claus S. Sørensen, Zoltan Szallasi

**Affiliations:** Biotech Research & Innovation Centre, University of Copenhagen, Copenhagen, Denmark; Translational Cancer Genomics Group, Danish Cancer Institute, Copenhagen, Denmark; Computational Health Informatics Program, Boston Children’s Hospital, Boston, MA; Department of Pathology, Brigham and Women’s Hospital, Boston, MA; Department of Medical Oncology, Dana-Farber Cancer Institute, Boston, MA; Department of Radiation Oncology, Dana-Farber Cancer Institute, Boston, MA; Harvard Medical School, Boston, MA; Department of Bioinformatics, Semmelweis University, Budapest, Hungary

**Keywords:** nucleotide excision repair, mutational signatures, cancer panel sequencing, urothelial cancer

## Abstract

Next generation sequencing based mutational signatures are frequently used to identify tumors with specific DNA repair deficiencies for targeted therapeutic strategies. Although mutational signatures are most commonly derived from whole exome (WES) or whole genome sequencing (WGS) data, more patients currently undergo tumor sequencing using more limited targeted panels that typically encompass several hundred cancer-associated genes. Identifying clinically relevant mutational signatures from targeted panel data requires new approaches capable of deriving signatures from the more limited sequencing data. Here, we derive and validate a panel sequencing-based composite mutational signature associated with nucleotide excision repair (NER) deficiency induced by inactivating *ERCC2* mutations in bladder cancer. Using publicly available panel sequencing data, we find that *ERCC2* wild type (WT) bladder cancer cases that have high levels of this mutational signature respond better to neoadjuvant platinum therapy and have improved overall survival compared to *ERCC2* WT cases with low levels of the signature. We also find that other solid tumor types with *ERCC2* mutations also show the characteristic mutational signature seen in NER-deficient *ERCC2*-mutant bladder cancers, suggesting a novel approach to therapeutically target these *ERCC2*-mutant solid tumors beyond bladder cancer.

## Introduction

Detecting specific DNA repair deficiencies of solid tumors in the clinical setting has significant implications for selecting the most effective therapy. Homologous recombination deficient tumors respond to PARP inhibitors (1), nucleotide excision repair (NER) deficient cancers respond to platinum based therapy (2,3), and mismatch repair deficient tumors are effectively treated with immune checkpoint inhibitors (4).

DNA repair deficiencies can be identified from the mutational profiles extracted from next generation sequencing data of tumor DNA. Mutational profiles are usually extracted from whole genome or whole exome sequencing data in the research setting (5,6), but routine clinical diagnostics has been mainly limited to capture-based targeted sequencing panels covering 400-500 genes spanning a few Megabases (Mb) of DNA. While these gene panels are designed to identify pathogenic mutations in known cancer-associated genes, they can also be exploited to derive DNA repair deficiency associated mutational signatures (7), albeit with lower accuracy and sensitivity than achievable with whole genome or whole exome data.

We previously reported that NER deficiency conferred by inactivating mutations in *ERCC2* is associated with a specific mutational profile in bladder tumors characterized by the COSMIC single base substitution (SBS) signature 5 and COSMIC indel (ID) signature 8 (5). We derived a composite mutational signature from bladder cancer WES datasets and showed that activity of this signature was not only associated with *ERCC2* inactivating mutations, but also identified likely NER deficient cases that lacked an *ERCC2* inactivating mutation but still exhibited increased sensitivity to platinum-based chemotherapy (5). Detecting NER deficiency is not only relevant for the identification of platinum sensitivity, but also opens the possibility for use of other NER deficiency specific drugs such as irofulven analogs (5,8,9). Given the existence of a large number of institutional and publicly available cases with available targeted panel sequencing and clinical outcomes data, we explored whether an accurate mutational signature of NER deficiency could be identified from targeted panel sequencing data.

## Methods

### Patients and cohorts

#### TCGA BLCA

The Cancer Genome Atlas (TCGA) Bladder Cancer (BLCA) whole-genome sequencing (WGS) cohort comprising 23 samples was used for mutation enrichment analysis. Normal and tumor WGS BAM files, aligned to the human reference genome (GRCh37/hg19), were downloaded from the GDC Data Portal (https://portal.gdc.cancer.gov/).

#### DFCI-MSKCC

The Dana-Farber Cancer Institute–Memorial Sloan Kettering Cancer Center (DFCI-MSKCC) cohort includes whole-exome sequencing data from pretreatment tumor and matched germline DNA from 50 patients with muscle-invasive or locally advanced urothelial carcinoma (3). All patients received cisplatin-based neoadjuvant chemotherapy (NAC) followed by cystectomy. Mutation data were obtained from cBioPortal (https://www.cbioportal.org/study/summary?id=blca_dfarber_mskcc_2014) (10). Normal and tumor BAM files were obtained from dbGaP (https://www.ncbi.nlm.nih.gov/gap/) under accession code phs000771.

#### Philadelphia

The Philadelphia cohort comprises whole-exome sequencing data from pre-chemotherapy tumor and matched germline DNA of 48 patients with muscle-invasive bladder cancer (MIBC) who received neoadjuvant chemotherapy (NAC) followed by cystectomy (11,12). Somatic single-nucleotide variants, identified with Mutect (13) and computationally filtered to remove artifacts from oxidative DNA damage during sample preparation (14), and short insertions/deletions (indels) were used in the analysis. Normal and tumor BAM files were obtained from dbGaP (https://www.ncbi.nlm.nih.gov/gap/) under accession code phs000771.

#### Aarhus

The Aarhus cohort includes 165 WES samples derived from patients with bladder cancer receiving chemotherapy (15). Of the 165 cases, 60 patients received NAC before cystectomy, and 105 patients received first-line chemotherapy upon detection of locally advanced or metastatic disease. Mutations from somatic VCF files after additional filtering were used in the analysis of the neoadjuvant subset (n=60).

#### GENIE

American Association for Cancer Research (AACR) GENIE data (v16.1-public) (16) was downloaded via the Synapse platform (https://synapse.org/genie). A small subset of samples from the GENIE dataset sequenced by the MSK-IMPACT panel with available clinical information (17) (n=38) that were not included in the training dataset were tested separately.

### Mutation enrichment analysis

In the TCGA BLCA WGS cohort, somatic single-base substitutions (SBS) and insertions/deletions (indels) were called using Mutect2 (GATK v3.8) (18). CallableLoci (GATK v3.8) was applied to identify genomic positions with sufficient read quality for mutation calling. The probability of observing a mutation in each sample was calculated as the total number of mutated bases divided by the total number of callable genomic positions. Enrichment or depletion of SBS and indels in specific genomic regions was determined by comparing the number of observed and expected mutations using two-sided binomial tests implemented in the MutationalPatterns R package (v3.18.0) (19). SBS and indel mutational spectra of tumor samples were constructed using the ICAMS (20) and MutationalPatterns R packages.

### Development of a prediction model

#### Training Dataset

The predictive model was trained on Bladder Urothelial Carcinoma (BLCA) samples from the GENIE dataset sequenced with the MSK-IMPACT panels (panel versions 341, 410, 468 and 505; n = 2077). Some samples originated from the same patient, which were either sequenced with a different panel version, at a different age of the patient, or the tumor type was different (primary vs. metastatic). To avoid introducing bias or overrepresenting individual patients in the analysis, we selected only one sample per patient, prioritizing (in order) those with primary tumor type, sequenced with a later version of the panel, or samples that had the highest number of mutations. We also filtered out samples with fewer than 5 mutations to exclude low-information profiles that could introduce noise and reduce the robustness of the training procedure. As a training set, we only chose samples that had either no *ERCC2* mutations (ERCC2-WT), or had a pathogenic *ERCC2* mutation (ERCC2-MUT). As pathogenic mutations, we considered the *ERCC2* variants which have been previously tested experimentally and determined to sensitize cells to cisplatin (2,17) (Supplemental Table 1). The final training set contained 1609 samples.

#### Test Datasets

The model was first validated on two independent subsets of the GENIE cohort (Supplemental Table 2): 1) BLCA samples with wild-type (WT) and known pathogenic *ERCC2* mutations (MUT) sequenced with the DFCI Oncopanel targeted panel sequencing platforms (panel versions 1, 2, or 3; n = 549), and 2) BLCA samples with WT and known pathogenic *ERCC2* mutations sequenced with any other targeted sequencing panel (n = 498, Supplemental Table 3). Second, we tested the model on three independent down-sampled WES datasets (DFCI-MSKCC, Aarhus, and Philadelphia) derived from patients with BLCA. The WES datasets were downsampled to the genes included in the latest version of the MSK-IMPACT panel (MSK-IMPACT 505). After downsampling to the MSK-IMPACT 505 panel, one sample from the DFCI-MSKCC cohort and two samples from the Aarhus cohort were excluded because no mutations were detected in the genes included in the MSK-IMPACT 505 panel. Finally, we tested the model on all experimentally validated, cisplatin-sensitizing pathogenic *ERCC2* variants (n = 44) found across various cancer types – excluding bladder cancer – and sequenced using any targeted panel platform within the GENIE cohort. Where applicable, the same filtering steps were applied to these datasets as the training dataset.

#### Features representing SBS and ID Mutational Signatures

For each dataset, single base substitution (SBS) and indel (ID) mutational spectra – comprising 96 and 83 mutation categories, respectively – were constructed using the MutationalPatterns (19) and ICAMS (20) R packages. These mutational spectra were then used to calculate the cosine similarity values between the mutational spectrum of each sample and the reference SBS and ID COSMIC signatures previously found to be present in bladder cancer (6).

#### Feature Harmonization and Selection with Recursive Feature Elimination

To identify the optimal subset of mutational features needed to train the panel-based ERCC2mut model, a recursive feature elimination (RFE) process using the caret R package (21) was performed on the training set. The input dataframe for RFE contained 18 features in total, including cosine similarities with SBS (SBS1, SBS2, SBS5, SBS8, SBS13, SBS29, SBS40) and ID (ID1, ID2, ID3, ID4, ID5, ID8, ID9, ID10) mutational signatures that have been previously identified in bladder cancer (6). The final three features were the harmonized (normalized and standardized) z-scores of single nucleotide variation (SNV) tumor mutational burden (TMB), deletion TMB, and insertion TMB (details are provided in the Supplemental Methods). Briefly, the TMB values were calculated as the total number of mutations of a certain type (SNV, DEL or INS) divided by the size of the panel. In order to make the values comparable across different panels, the TMB values were then normalized using the ordered quantile normalizing transformation function from the bestNormalize R package (22), followed by standardization to z-scores (Supplemental Figure 1).

In the subset of pathogenic *ERCC2* variants from other cancer types, we performed a panel- and cancer type-specific harmonization to account for differences in TMB not just across sequencing platforms but also across cancer types.

Model performance at each step of RFE was evaluated using the area under the ROC curve (AUC), along with sensitivity (true positive rate) and specificity (true negative rate). To ensure robustness of the feature selection process, RFE was performed with 5-fold cross-validation (CV). RFE was then carried out with a gradient boosting model (gbm) as the classification algorithm. Candidate feature subsets ranging from 2 to 10 predictors were evaluated. The selection metric was the ROC AUC, which was maximized to identify the optimal subset of features.

#### Training Procedure of a gradient boosted model classifier

A gradient boosted model (GBM) classifier was trained to predict *ERCC2* mutation status. The training procedure consisted of three steps: 1) feature selection and preprocessing, 2) hyperparameter tuning, and 3) model training. Further details are provided in the Supplemental Materials.

#### Optimal cutpoint

To determine the optimal threshold for classifying *ERCC2* mutation status, the cutpointr R package was used (23). The function evaluates a range of possible thresholds and identifies the one that maximizes classification performance according to the default criterion (sum of sensitivity and specificity). This optimal cutpoint was extracted and used to convert predicted probabilities into binary class assignments, such that samples that had prediction scores higher than the threshold were classified as ERCC2-MUT. Optimal cut-off values were determined independently for each of the two testing sets to maximize accuracy and other performance metrics (Supplemental Table 4).

### Survival analysis

Survival curves were estimated using the Kaplan-Meier method with the survival and survminer R packages. The Cox proportional hazards (CPH) model was used to model the effects of multiple covariates on the survival time, including the panel-based ERCC2mut score (as binary variable), the patient’s age at the time the sequencing result was reported, and the patient’s race. Samples were grouped by sample type according to their origin, either primary or metastatic. The proportional hazards assumption was fulfilled for all covariates. Survival time was calculated as the difference in days between the date of death or last contact and the patient’s age in days at the time the sequencing result was reported.

### Data availability

The Cancer Genome Atlas (TCGA) Bladder Cancer (BLCA) whole-genome sequencing (WGS) normal and tumor BAM files, aligned to the human reference genome (GRCh37/hg19), were downloaded from the GDC Data Portal (https://portal.gdc.cancer.gov/). Whole-exome sequencing (WES) normal and tumor BAM files from the Dana-Farber Cancer Institute–Memorial Sloan Kettering Cancer Center (DFCI-MSKCC) cohort were obtained from dbGaP (https://www.ncbi.nlm.nih.gov/gap/) under accession code phs000771. Mutation data were obtained from cBioPortal (https://www.cbioportal.org/study/summary?id=blca_dfarber_mskcc_2014). Philadelphia WES normal and tumor BAM files are also available through dbGaP under accession code phs000771. Raw sequencing files from the Aarhus cohort are available at https://www.ebi.ac.uk/ega/studies/EGAS00001004507. Data from the American Association for Cancer Research (AACR) GENIE project (v16.1-public) were downloaded via the Synapse platform (https://synapse.org/genie).

### Code availability

There are no restrictions to access the custom code used for the analyses presented in this study. Information is available from the authors on request. The code implementing the panel-based ERCC2mut predictive model is publicly available on GitHub at https://github.com/jud-b/ERCC2mut.

## Results

### The mutational patterns present in *ERCC2*-mutant tumors are enriched in genomic regions covered by targeted sequencing panels

Transcriptionally active regions may be particularly vulnerable to mutations due to transcription-associated mutagenesis (24). This increased mutagenic rate can be counterbalanced in part by transcription-coupled nucleotide excision repair (TC-NER) (25). Since *ERCC2* plays a critical role in TC-NER, and genes included in targeted seqencing panels are often highly transcribed, we hypothesized that panel sequencing data are enriched for mutations arising in the setting of NER deficiency conferred by pathogenic *ERCC2* mutations (17). Using the TCGA bladder cancer (BLCA) whole-genome sequencing (WGS) data, we calculated the number of single base pair substitution (SBS) and insertion/deletion (ID) mutations for *ERCC2* mutant and wild type bladder cancer cases at regions of the genome with a sufficient number of high quality reads to call mutations. These total mutation counts were used to calculate the observed and expected number of mutations in the genomic regions contained on the various panel sequencing platforms (Supplemental Table 5). In these genomic regions, we found a significant enrichment in the total number of SBS mutations (Figure 1A, two-sided binomial tests with multiple testing correction) in *ERCC2* MUT tumors. This enrichment was especially marked for T>C mutations, the mutation type dominating SBS5, a mutation signature associated with *ERCC2* mutations (Figure 1B, pairwise Wilcoxon rank-sum tests). Additionally, the observed number of indels in panel sequencing regions in *ERCC2*-mutant samples was significantly higher than expected (Figure 1C, two-sided binomial tests with multiple testing correction). Furthermore, the increased number of indels was dominated by >5 bp deletions, which is a hallmark of the ID8 signature (Figure 1D, pairwise Wilcoxon rank-sum tests) that we previously identified as significantly associated with the presence of *ERCC2* inactivating mutations (5). Interestingly, the observed number of both SBS mutations and indels were lower than expected in the panel sequencing data of *ERCC2* WT cases (Figure 1A and C), suggesting that transcription-coupled repair (TCR) is effectively removing these types of mutations in NER proficient cancer cells. Taken together, these results suggest that the number of *ERCC2* mutation associated SBS and short indel mutations are enriched in panel sequencing data relative to their expected background frequencies and that *ERCC2* inactivation associated mutational events may therefore be detected with sufficient frequency from panel sequencing data to warrant further investigation as a potential biomarker.

**Figure 1:**
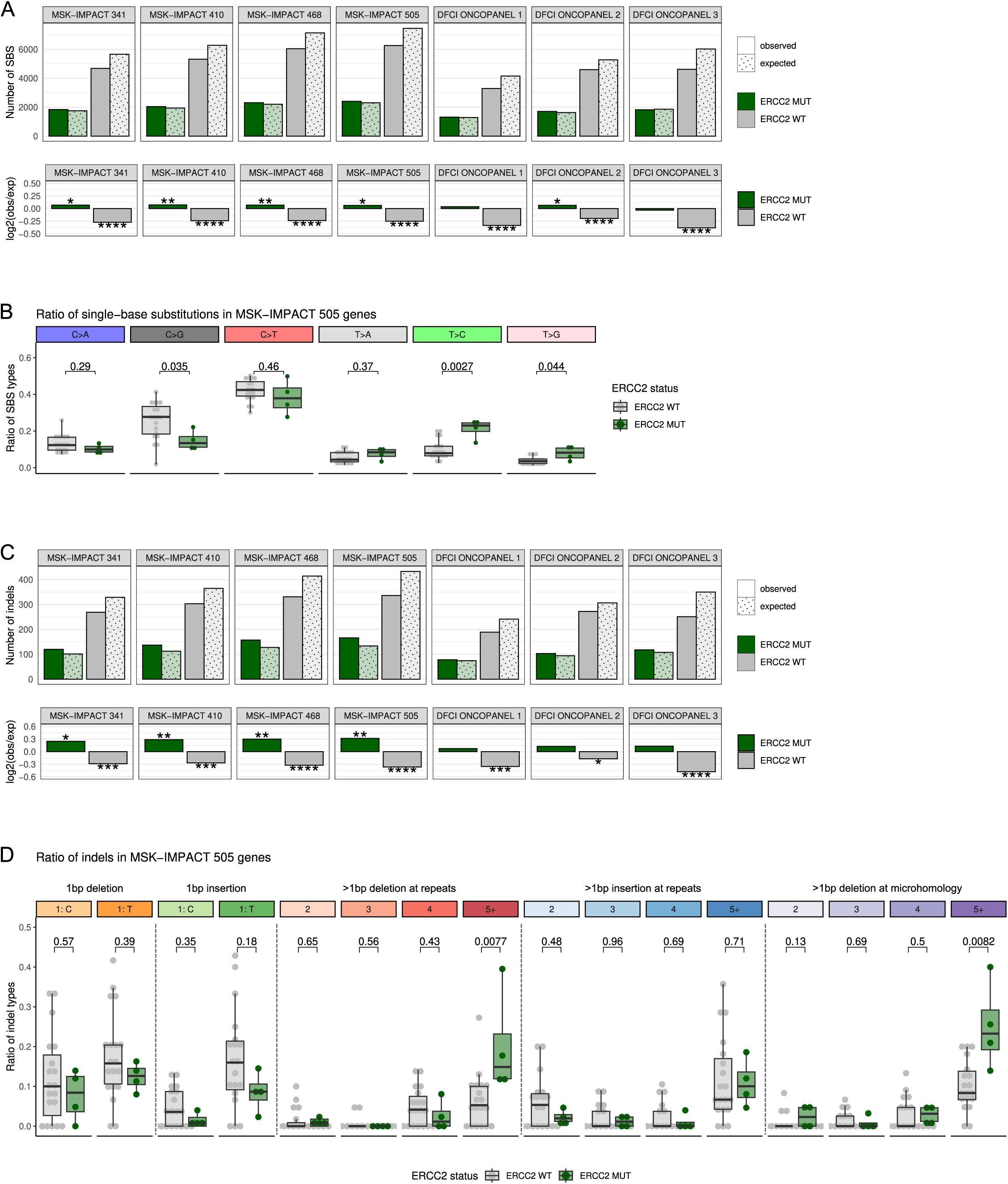
Enrichment of single-base substitutions (SBS) and indels in genomic regions covered by targeted sequencing panels. **A.** Observed versus expected number of SBS, shown as counts and log ratios in genes included in the MSK-IMPACT (panel versions 341, 410, 468, 505) and DFCI Oncopanel (panel versions 1, 2, 3) targeted sequencing panels. Enrichment or depletion of mutations in the defined genomic regions was tested using two-sided binomial tests with multiple testing correction setting the false discovery rate (FDR) at α = 0.1. **B.** Ratio of major SBS types in *ERCC2*-mutant (MUT) versus wild-type (WT) samples (pairwise Wilcoxon rank-sum tests). **C.** Observed versus expected number of indels, shown as counts and log ratios in MSK-IMPACT and DFCI Oncopanel genes. Enrichment or depletion of mutations in the defined genomic regions was tested using two-sided binomial tests with multiple testing correction setting the false discovery rate (FDR) at α = 0.1. **D.** Ratio of major indel types in *ERCC2* MUT versus WT samples (pairwise Wilcoxon rank-sum tests). The colors along the x axis correspond to the Catalog of Somatic Mutations in Cancer (COSMIC) indel categories: 1 bp deletions with the deleted base indicated (C or T), 1 bp insertions with the inserted base indicated (C or T), >1bp deletions at repeats with the deletion length indicated, >1bp insertions at repeats with the insertion length indicated, and >1bp deletions at microhomologies with the deletion length indicated. Significance levels of FDR adjusted p-values: *P ≤ 0.1, **P ≤ 0.05, ***P ≤ 0.01, ****P ≤ 0.001.

### A panel-based composite mutational signature of *ERCC2* inactivating mutations

The enrichment of *ERCC2*-inctivation associated mutations in panel-sequenced regions suggested that a panel-based classifier was feasible. We therefore developed a gradient boosting machine (GBM) classifier (from now on “panel-based ERCC2mut signature”) that accurately differentiates between *ERCC2* MUT and WT cases (Figure 2A). The model was trained using bladder urothelial carcinoma (BLCA) samples sequenced with the MSK-IMPACT targeted sequencing panels (n = 1609) within the GENIE dataset (16). Features were selected from the SBS and ID mutational signatures previously identified in bladder cancer sequencing data (5,26). In addition, the total number of SBS and ID mutations in the panel sequencing data were also considered as potential discriminating features (see Methods). Six features – SBS signatures 2, 5, and 13, ID signature 8, and the total number of SBS and ID mutations – were combined in the final GBM classifier (Figure 2B,C). The optimal threshold of >0.048 for classifying samples as *ERCC2* MUT was determined by maximizing the sum of sensitivity and specificity values (Supplemental Table 4). The final panel-based ERCC2mut signature achieved good performance metrics in the training dataset including 89.4% accuracy, 98.2% sensitivity, 88.8% specificity (Table 1), and demonstrated an excellent ability to distinguish between the *ERCC2* MUT and WT cases with an area under the curve (AUC) of 0.98 (Figure 2D,E; Supplemental Table 6).

**Figure 2:**
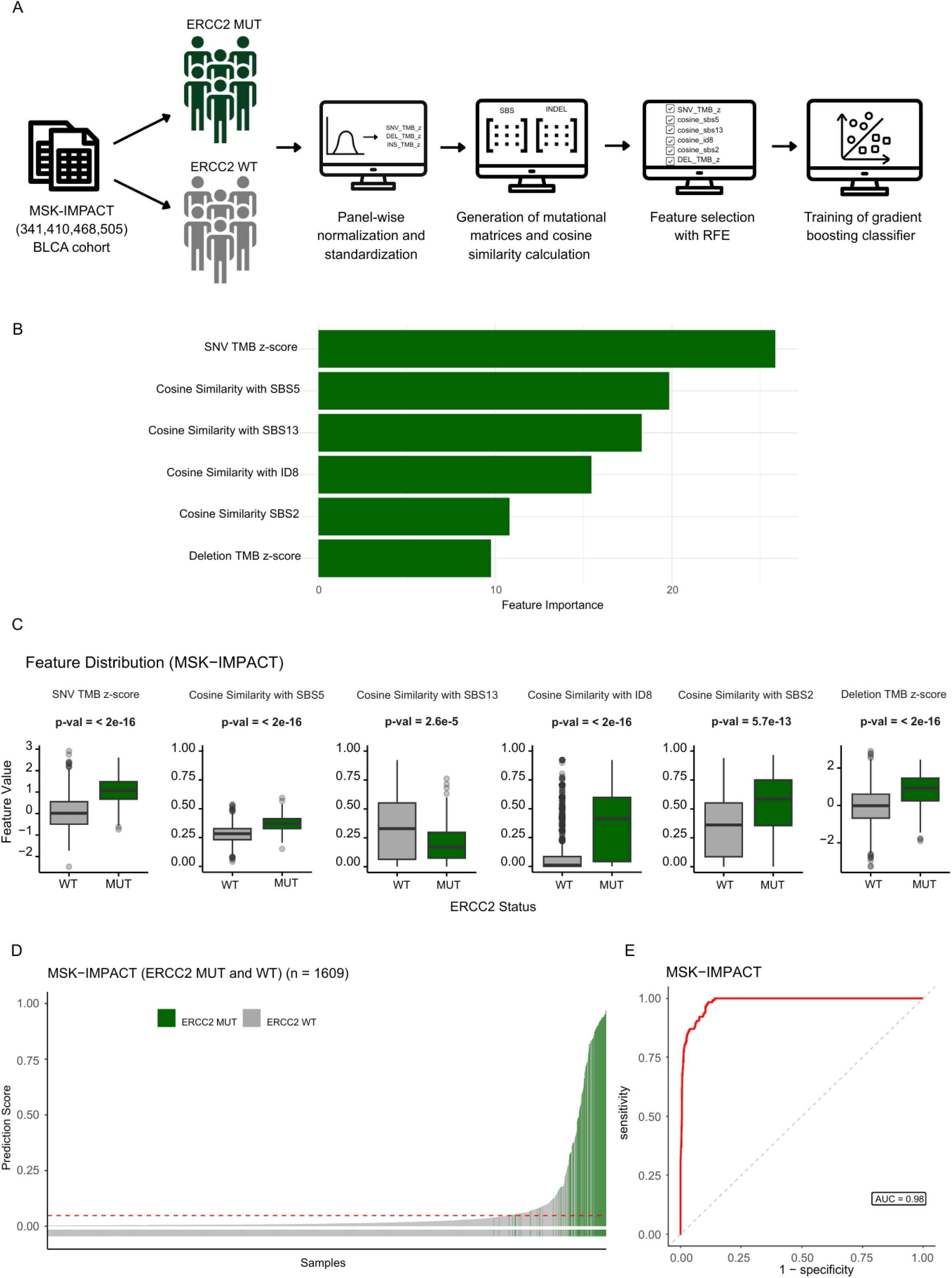
Development of a panel-based gradient boosting machine (GBM) classifier of *ERCC2* mutation status. **A.** Schematic representation of the model-building workflow. **B.** Importance scores of the selected features in the panel-based ERCC2mut BLCA model. **C.** Feature distribution in the training set of the panel-based ERCC2mut classifier. P-values were calculated using the Wilcoxcon rank-sum test. **D.** Distribution of prediction scores for the training dataset (MSK-IMPACT). **E.** Receiver operating characteristic (ROC) curve and area under the curve (AUC) value for the training dataset (MSK-IMPACT).

**Table 1:** Evaluation metrics and cut-off values for the panel-based ERCC2mut classifier in training and validation datasets. Summary of classifier performance in the training dataset (MSK-IMPACT) and in two independent validation datasets within the GENIE cohort (the DFCI-Oncopanel and the pooled dataset of 14 additonal panels with BLCA samples). Reported metrics include accuracy, sensitivity, specificity, positive predictive value (PPV), negative predictive value (NPV), and balanced accuracy, along with the optimal cut-off values used for classification.

|  | Cut-off | Accuracy | Sensitivity | Specificity | Positive Predictive Value (PPV) | Negative Predictive Value (NPV) | Balanced accuracy |
| --- | --- | --- | --- | --- | --- | --- | --- |
| MSK-IMPACT | 0.04821027 | 89.4% | 98.2% | 88.8% | 40% | 99.9% | 93.5% |
| DFCI-ONCOPANEL | 0.1279685 | 91.6% | 84.6% | 92% | 34.4% | 99.2% | 88.3% |
| Pooled panels | 0.1322924 | 88.5% | 81.8% | 88.8% | 25% | 99.1% | 85.3% |

Next, we tested the panel-based ERCC2mut predictor – trained on the MSK-IMPACT GENIE dataset – on the DFCI-ONCOPANEL GENIE dataset (549 cases). The predictor accurately identified the *ERCC2*-mutant cases, achieving an AUC of 0.9 (Figure 3 A, B), 91.6% accuracy, 84.6% sensitivity, 92% specificity (Table 1), and correctly classifying 22 out of 26 *ERCC2*-mutant cases (Supplemental Table 7). In addition, another 42 cases exhibited the *ERCC2* mutation-associated composite mutational signature without the presence of a known pathogenic *ERCC2* mutation (Supplemental Table 7). We then pooled together all non-MSK-IMPACT and non-DFCI-Oncopanel cohorts of BLCA samples from the GENIE database and tested the performance of the predictor and attained similar results: AUC of 0.9, 88.5% accuracy, 81.8% sensitivity, and 88.8% specificity (Figure 3C,D; Table 1). 18 out of 22 *ERCC2*-mutant samples were identified, with an additional 53 *ERCC2* wild-type cases exhibiting the mutational signature (Supplemental Table 8).

**Figure 3:**
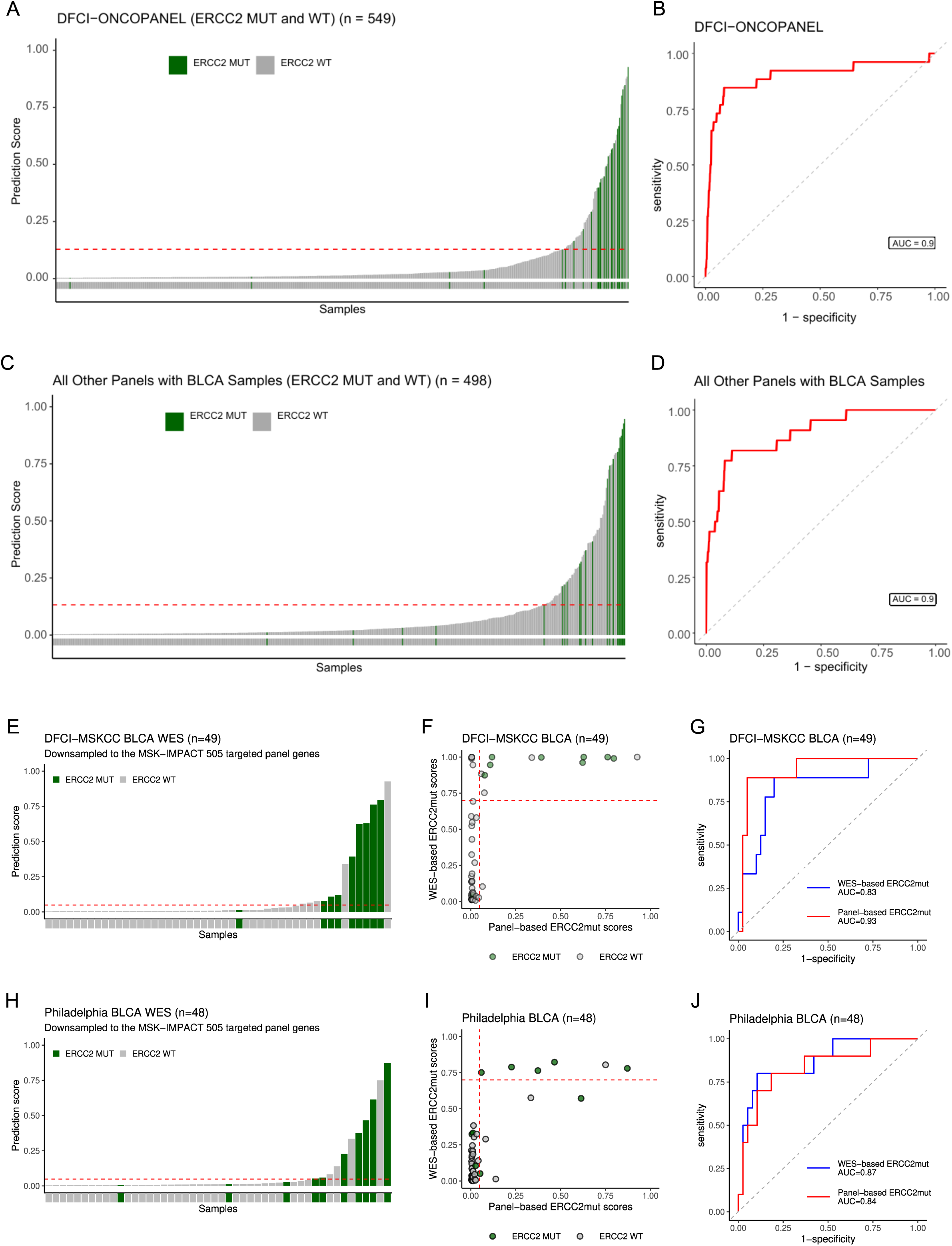
Comparison of panel-based and WES-based ERCC2mut models and performance of the panel-based ERCC2mut classifier on the test datasets. **A.** Distribution of prediction scores for the DFCI-Oncopanel test dataset. **B.** ROC curve and AUC value for the DFCI-Oncopanel dataset. **C.** Distribution of prediction scores for the pooled test dataset assembled from 14 targeted panels with BLCA samples from the GENIE dataset. **D.** ROC curve and AUC value for the pooled test dataset. **E.** Distribution of panel-based ERCC2mut scores in the DFCI-MSKCC WES cohort calculated using mutations downsampled to genes present in the MSK-IMPACT 505 panel. **F.** Correlation between the panel-based and WES-based ERCC2mut scores in the DFCI-MSKCC cohort. The red dashed lines indicate the cut-off values used to separate *ERCC2* MUT and WT cases: 0.048 for the panel-based model (x axis) and 0.7 for the WES-based model (y axis). **G.** ROC curve and AUC for the panel-based and WES-based ERCC2mut models in the DFCI-MSKCC cohort. **H.** Distribution of panel-based ERCC2mut scores in the Philadelphia WES cohort downsampled to genes present in the MSK-IMPACT 505 panel. **I.** Correlation between the panel-based and WES-based ERCC2mut scores in the Philadelphia cohort (Cohen’s Kappa = 0.649, 1.6 10^-6^). The red dashed lines indicate the cut-off values used to separate *ERCC2* MUT and WT cases: 0.048 for the panel-based model (x axis) and 0.7 for the WES-based model (y axis). **J.** ROC curve and AUC for the panel-based and WES-based ERCC2mut models in the Philadelphia cohort.

### The panel-based ERCC2mut signature is associated with *ERCC2* mutant bladder cancer cases

We previously found that bladder cancer cases harboring the WES-based *ERCC2* mutational signature respond to platinum therapy, even in the absence of an inactivating *ERCC2* mutation (5). If a similar predictor could be derived from panel sequencing data, it may allow the evaluation of large bladder cancer cohorts where such data are available. To investigate this possibility, we used bladder cancer whole exome sequencing data from previously published cohorts: DFCI-MSKCC (3), Philadelphia (11,12), and Aarhus (15), which we downsampled to the regions matching the MSK-IMPACT gene panel’s 505 genes. In the DFCI-MSKCC cohort, we previously calculated a WES-derived *ERCC2* mutation-associated composite mutational signature score (5). Cases with a score >0.7 were considered to be NER deficient and showed platinum sensitivity even in the absence of inactivating *ERCC2* mutations. We calculated the panel-based ERCC2mut signature scores for the DFCI-MSKCC cases using only downsampled WES data corresponding to the exonic regions covered by the panel (Figure 3E). We assessed whether cases with >0.7 *ERCC2* inactivation-associated mutational signature scores derived from WES also have higher panel-based ERCC2mut signature scores (details regarding the WES- and panel-based ERCC2mut models are provided in Supplemental Table 9). Indeed, we found substantial agreement between the original WES-based and the downsampled panel-based *ERCC2*mut signature activities (Figure 3F, Cohen’s Kappa = 0.714, p-value = 3.6·10^-7^). Out of 17 cases with the *ERCC2* mutational signature based on WES data, 13 were also predicted to be *ERCC2*-mutant in the downsampled panel sequencing data (i. e., value >0.048), with eight of those harboring known pathogenic *ERCC2* mutations (Supplemental Table 10). Both the panel-based and the WES-based models identified all true pathogenic *ERCC2*-mutant samples except one and distinguished between *ERCC2* WT and *ERCC2* MUT classes (Figure 3G, AUC_panel_ = 0.93 and AUC_WES_ = 0.83). This result suggests that some, but not all, cases displaying the *ERCC2* inactivation associated composite mutational signature in the WES data can also be detected by panel sequencing data. Similar outcomes were observed with the Philadelphia (Figure 3H-J, Supplemental Table 11) and Aarhus (Supplemental Figure 2, Supplemental Table 12) WES bladder cancer cohorts.

### The panel-based ERCC2mut signature is associated with response to neoadjuvant chemotherapy and improved survival in the absence of inactivating *ERCC2* mutations

Patients in the DFCI-MSKCC, Philadelphia, and Aarhus cohorts are comprised of muscle-invasive bladder cancer (MIBC) patients who received neoadjuvant cisplatin-based chemotherapy (NAC) followed by radical cystectomy. These cases together with a subset of neoadjuvant cases with available clinical information (17) from the GENIE cohort sequenced by the MSK-IMPACT panel were combined for a pooled analysis (Supplemental Table 13). Cisplatin responders were defined as those patients with pathologic downstaging of tumors to nonmuscle invasive, node-negative disease (i.e., pT0, pTa, pTis, or pT1; and N0) at the time of cystectomy, whereas nonresponders were patients with residual muscle-invasive (pT2) or node-positive (≥N1) disease. In this combined neoadjuvant cohort, the panel-based ERCC2mut signature was associated with response to neoadjuvant platinum-based therapy even in the absence of pathogenic *ERCC2* mutations (Figure 4A,B; Fisher’s exact test: p-value = 0.021). This association strengthened if a stricter definition of response (pT0, pTa, or pTis; and N0) was applied (Supplemental Figure 3, Fisher’s exact test: p-value = 0.01).

**Figure 4:**
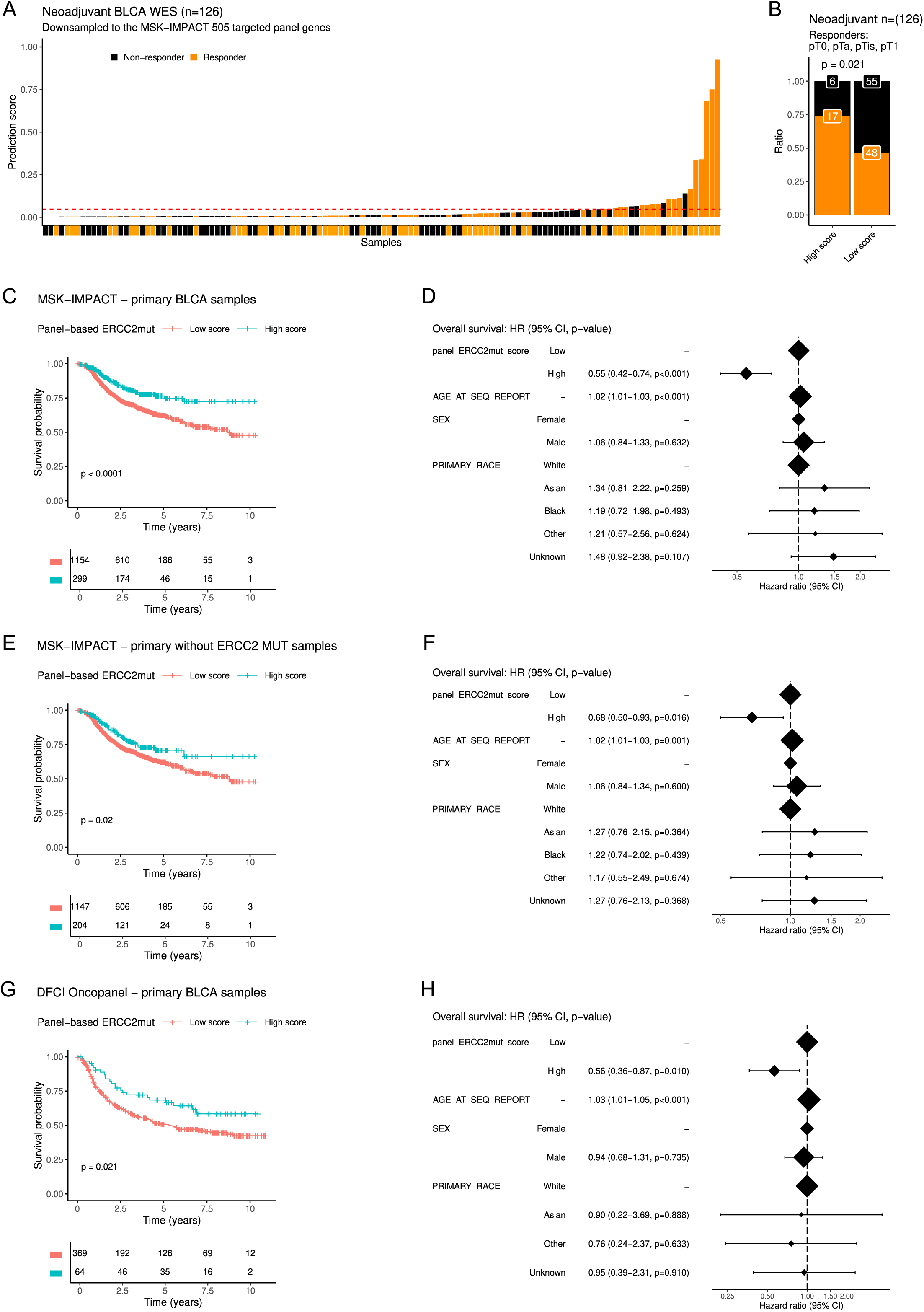
Panel-based ERCC2mut scores in MIBC, survival curves estimated by the Kaplan-Meier method and hazard ratios estimated by the Cox Proportional Hazards (CPH) model. **A.** Distribution of panel-based ERCC2mut scores in the combined neoadjuvant MIBC cohort excluding pathogenic *ERCC2* MUT cases (only *ERCC2* WT cases) colored by response to neoadjuvant platinum-based chemotherapy. **B.** Bar plots showing the ratio of responders and nonresponders among *ERCC2* WT cases with high and low panel-based ERCC2mut scores (threshold value = 0.048) in the neoadjuvant MIBC cohort excluding pathogenic *ERCC2* MUT cases (Fisher’s exact test: p-value = 0.021). Response was defined as pT0, pTa, pTis, and pT1. **C.** Kaplan-Meier survival curves and risk table for all BLCA cases with primary samples sequenced in the MSK-IMPACT dataset, stratified by the panel-based ERCC2mut prediction score as binary classes (cut-off: 0.048; log-rank test: p-value < 0.0001). **D.** Hazard ratios with 95% confidence intervals (CIs) and p-values estimated by the CPH model. **E.** Kaplan-Meier survival curves and risk table for BLCA cases with primary samples sequenced in the MSK-IMPACT dataset, excluding known pathogenic *ERCC2* mutations, stratified by the panel-based ERCC2mut prediction score (cut-off: 0.048; log-rank test: p-value = 0.02). **F.** Hazard ratios with 95% confidence intervals (CIs) and p-values estimated by the CPH model. **G.** Kaplan-Meier survival curves and risk table for all BLCA cases in the DFCI-Oncopanel dataset, stratified by the panel-based ERCC2mut prediction score (cut-off: 0.128; log-rank test: p-value = 0.021). **H.** Hazard ratios with 95% confidence intervals (CIs) and p-values estimated by the CPH model.

We next investigated whether the presence of the panel-based ERCC2mut signature correlates with improved survival for patients with bladder cancer in the panel sequencing cohorts from the GENIE dataset. Unfortunately, patient age, tumor stage at diagnosis, and treatment history were not publicly available. To approximate overall survival, we calculated the time between the patient’s age at the sequencing report and their age at last contact or death, acknowledging that the unknown interval between diagnosis and sequencing is a potential source of bias. Patients were analyzed separately based on whether the sequenced sample was from the primary tumor or a metastatic site. In the MSK-IMPACT dataset of primary samples, patients with a high panel-based ERCC2mut signature demonstrated significantly better survival (Figure 4C, Log-rank test: p < 0.0001) and lower risk of death (Figure 4D, HR = 0.55, 95% CI 0.42–0.74, p<0.001) compared to cases with low panel-based ERCC2mut scores. This association remained significant even after excluding cases with known pathogenic *ERCC2* mutations (Figure 4E, Log-rank test: p = 0.02 and Figure F, HR = 0.68, 95% CI 0.50-0.93, p = 0.016), suggesting that the panel-based ERCC2mut signature score may serve as a correlate of survival even in the absence of an *ERCC2* mutation. A similar trend was observed in the DFCI-Oncopanel dataset of primary samples (Figure 4G, Log-rank test: p=0.021 and Figure 4H, HR = 0.56, 95% CI 0.36-0.87, p = 0.01); however, the association did not remain significant in this cohort after the cases with known pathogenic *ERCC2* mutations were removed (Supplemental Figure 4A, Log-rank test: p=0.19 and Supplemental Figure 4B, HR = 0.71, 95% CI 0.44-1.12, p = 0.143). In the datasets of metastatic samples, we did not observe significant differences in OS based on panel-based ERCC2mut signature scores (Supplemental Figure 5).

### The panel-based ERCC2mut signature in *ERCC2* mutant cases beyond bladder cancer

Inactivating *ERCC2* helicase domain mutations are present almost exclusively in urothelial tumors. However, in rare instances, the most frequent inactivating mutations of *ERCC2* observed in bladder tumors (e.g. N238S, S44L, T484M) are also observed in other cancer types (Figure 5A). We had access to panel sequencing data for 55 such non-bladder *ERCC2* mutant cases, and 44 of these cases passed the filtering criteria (>5 mutations per sample and only including one sample per patient). The panel-based ERCC2mut signature was present in 31/44 (70%) of these cases (Figure 5B). Interestingly, some of these cases, especially those annotated as ‘cancer of unknown primary’, may be metastases from bladder cancer. However, others, such as the breast cancer case, are likely to be genuine non-bladder cancer cases. This would indicate that *ERCC2* inactivation leads to NER deficiency and possibly to platinum sensitivity in rare instances when present in non-bladder cancer cases.

**Figure 5:**
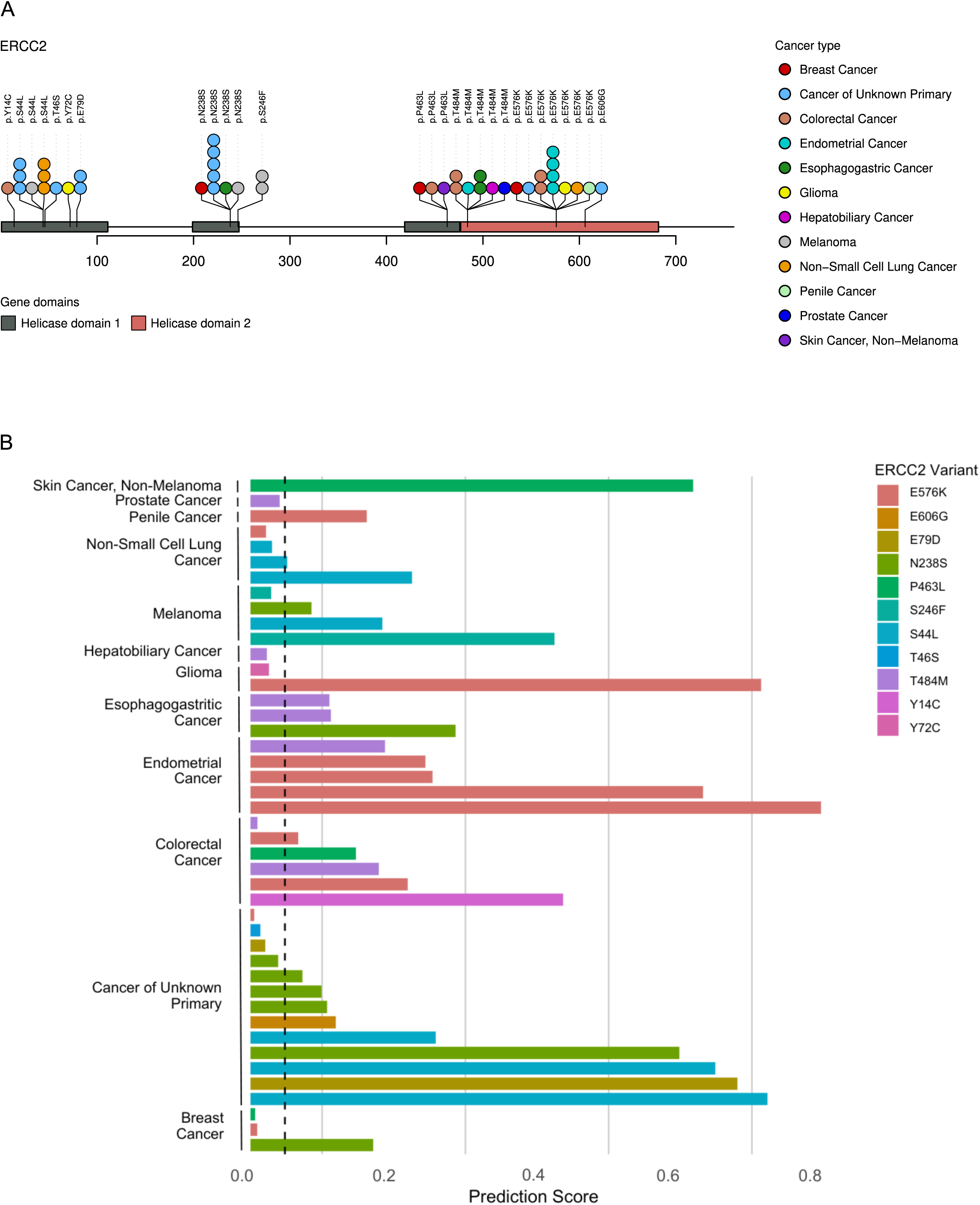
ERCC2-mutant cancer cases beyond bladder cancer. **A.** Location of experimentally verified pathogenic cisplatin-sensitive *ERCC2* mutations along the gene domains found in other cancer types. **B.** Panel-based ERCC2mut scores of these *ERCC2*-mutant cases indicating the presence of the composite mutational signature and potential repair deficiency. The black dashed line marks the cut-off value (0.048) used to classify samples as NER proficient or NER deficient.

## Discussion

Next generation sequencing based mutational signatures can identify DNA repair deficient cancer cases beyond those that are caused by inactivating mutations of individual DNA repair genes. For example, homologous recombination (HR) deficient cases without a loss of function mutation in a canonical HR gene can be identified by next generation sequencing based strategies (27,28), and some of those diagnostic methods have significant clinical utility for the prioritization of patients for PARP inhibitor therapy (29,30). Similarly, microsatellite unstable tumors can also be identified by specific mutational profiles (6), and such cases without an inactivating mismatch repair gene mutation also benefit from immune checkpoint inhibitor therapy (31).

We previously investigated whether we can identify NER deficient tumors that lack an obvious inactivating mutation of a key NER gene such as *ERCC2*. We derived a composite mutational signature from whole exome sequencing (WES) data from *ERCC2*-mutant cases (5), and we showed that tumors that lacked an *ERCC2* mutation but exhibited the mutational signature were sensitive to platinum treatment, similar to those cases with an *ERCC2* mutation (5). Because NER-deficient tumors are sensitive to platinum agents and irofulven analogs (17), a diagnosic tool for identifying NER deficient cancer cases could inform use of NER targeted therapies.

The accuracy of DNA repair deficiency associated mutational signature detection increases with the extent of genomic regions covered by NGS (27). Therefore, WES or WGS data are preferred to derive DNA repair deficiency specific mutational signatures. However, such data are often available only in specific research settings and not as part of routine clinical practice. While panel sequencing was introduced to clinical practice in the mid 2010s (32), routine whole exome sequencing is still at its earliest stages (33). This discrepancy motivated us to develop a panel sequencing based composite mutational signature of NER deficiency that could allow other researchers to mine panel sequencing data available at clinical centers.

We found that short deletions (ID8) and other mutational features associated with NER deficiency are enriched in the genomic regions represented in routine sequencing panels, thus allowing us to develop a panel sequencing based composite mutational signature that identified *ERCC2* mutant cases with similar accuracy to that seen in WES based cohorts. We showed that the presence of this mutational signature is associated with better survival in both *ERCC2* mutant and *ERCC2* WT subsets of patients. A major limitation of our analysis is the lack of detailed clinical information regarding treatment and outcomes. Nevertheless, the composite mutational signature developed here offers a framework for performing similar analyses at clinical centers with significant bladder cancer patient populations.

*ERCC2* mutations are rare in solid tumor types other than bladder cancer. Nevertheless, demonstrating that such cases frequently have a similar mutational signature as *ERCC2* mutant bladder cancer suggests that such cases are also NER deficient and may be sensitive to platinum based therapy. This could be useful information, particularly if platinum is not a standard therapy for the particular cancer type.

The panel sequencing based mutational signature presented here may be useful in further investigating the connection between NER deficiency and response to neoadjuvant platinum-based chemotherapy. Given the high pathologic complete response rates observed in bladder cancer patients with tumor DNA repair gene alterations, several clinical trials such as Alliance A031701 (NCT03609216) are investigating whether patients with localized bladder cancer and good response to chemotherapy could safely forego radical cystectomy. Although NER deficiency conferred by inactivating mutations in the helicase domains of *ERCC2* is the most common mechanism that renders bladder tumors sensitive to platinum, other mechanisms of NER deficiency may exist and could be identified by the reevaluation of the large amount of available panel sequencing data.

## Additional information

## Author contributions

OS, JB, CSS, KWM, and ZS conceptualized the project. OS and JB developed the methodology. ZsS, AP, MD provided computational resources. OS and JB performed the analysis. OS and JB were responsible for visualization. AN provided clinical information and insight. CSS, ZS, and KWM supervised the project. OS, JB, CSS, KM, and ZS wrote the original draft of the manuscript. OS, JB, ZsS, AP, MD, AN, CSS, KWM, and ZS edited the manuscript.

### Acknowledgments

Results shown here are based in part from data generated by the TCGA Research Network: https://www.cancer.gov/ccg/research/genome-sequencing/tcga and the International Cancer Genome Consortium (ICGC). Results presented in the current publication are based, in part, on the use of study data downloaded from the dbGaP website, under phs000771.v2.p1 accession code (https://www.ncbi.nlm.nih.gov/projects/gap/cgi-bin/study.cgi?study_id=phs000771.v2.p1). The authors would like to acknowledge the American Association for Cancer Research and its financial and material support in the development of the AACR Project GENIE registry, as well as members of the consortium for their commitment to data sharing. Interpretations are the responsibility of study authors.

