## Supplemental Materials for "Identification of an *ERCC2* mutation associated mutational signature of nucleotide excision repair deficiency in targeted panel sequencing data"

### 1 Supplemental Methods

#### 1.1 Harmonization of Tumor Mutational Burden (TMB)

Tumor mutational burden (TMB) values were calculated separately for single nucleotide variants (SNV), insertions (INS) and deletions (DEL) as the number of mutations per megabase (Mb) of sequenced region. The size of each sequencing panel was determined by calculating the total number of targeted bases expressed in megabases, to ensure that mutation counts were normalized by panel coverage.

To enable comparison across panels, TMB values were normalized and standardized within each panel. Normalization was performed using the ordered quantile normalizing transformation from the *bestNormalize* R package [1] to approximate a normal distribution of the TMB values. A small random jitter was added prior to transformation to prevent rank ties. Standardization was then performed by converting the transformed TMB values into z-scores. The TMB z-scores were calculated as the difference between the transformed TMB value and the respective panel-specific mean divided by the standard deviation.

For the validation dataset containing samples with pathogenic *ERCC2* mutations from other cancers, normalization and standardization were performed with respect to each cancer type and panel, accounting for potential differences in mutational landscapes across tumor types.

#### 1.2 Feature Selection and Preprocessing

A set of six features, determined by RFE (SNV_TMB_z, cosine_sbs5, cosine_sbs13, cosine_id8, cosine_sbs2, DEL_TMB_z) was used as predictors in the BLCA model, with *ERCC2* mutation status (ERCC2-MUT and WT) as the response variable. The same six features were selected for both the BLCA and the UTUC model. Missing values were imputed with zero, and the response variable was encoded as a binary outcome (ERCC2-MUT = 1, ERCC2-WT = 0).

#### 1.3 Hyperparameter tuning

To determine the optimal model configuration, a grid search was conducted across a broad range of hyperparameters, including the number of trees (50–50,000 in steps of 50), interaction depth (3, 5, 7, 9), learning rate (0.00001 to 0.1), and minimum node size (10). A 5-fold CV was applied using the *caret* R package [2, 3], and the parameter set yielding the best performance was selected for model training.

| Panel | Size (Mb) |
|---|---|
| MSK-IMPACT341 | 0.926688 |
| MSK-IMPACT410 | 1.049993 |
| MSK-IMPACT468 | 1.164911 |
| MSK-IMPACT505 | 1.254333 |
| DFCI-ONCOPANEL-V1 | 1.237438 |
| DFCI-ONCOPANEL-V2 | 1.422901 |
| DFCI-ONCOPANEL-V3 | 1.319975 |
| COLU-CCCP-V1 | 1.26216 |
| MDA-409-V1 | 1.749576 |
| PROV-TSO500HT-V2 | 1.970825 |
| UCSF-IDTV5-TN | 5.246771 |
| UCSF-IDTV5-TO | 5.246771 |
| UCSF-NIMV4-TN | 1.374256 |
| UCSF-NIMV4-TO | 1.374256 |
| UHN-TSO500-V1 | 1.954831 |
| UHN-555-V1 | 1.601237 |
| VICC-01-T7 | 1.565857 |
| VICC-02-XTV3 | 3.711446 |
| VICC-02-XTV4 | 3.711446 |
| VHIO-OCP-V3 | 40.12477 |
| YALE-OCP-V3 | 0.396357 |

**Supplemental Methods Table 1:** Targeted panels from the GENIE cohort with bladder urothelial carcinoma (BLCA) samples and their respective base coverage size (Mb).

| Hyperparameter | Value |
|---|---|
| Number of trees | 700 |
| Interaction depth | 9 |
| Learning rate | 0.005 |
| Bagging fraction | 0.2 |
| Node size | 10 |

**Supplemental Methods Table 2:** Hyperparameter values for training the panel-based ERCC2mut classifier.

#### 1.4 Model training

Using the tuned hyperparameters, two GBM models were trained under a Bernoulli distribution with a bagging fraction of 0.2. For the training of the models, we used the *gbm3* R package [4]. A two-step training procedure was applied, both steps using the training set that consists of ERCC2-WT and ERCC2-MUT BLCA samples sequenced with the MSK-IMPACT panels. In the first step, the original training set was stratified with respect to the two classes (ERCC2-WT and ERCC2-MUT) and a model was trained on 80% of the data while it was validated on the remaining 20%, using the tuned hyperparameters. The final model was then retrained on the full training dataset using the same hyperparameters.

##### 1.5 Evaluation metrics

The performance of the classifier was evaluated by comparing predicted class labels against the true *ERCC2* mutation status. Several key performance metrics were used to assess the performance of the model:

- **Accuracy:** the proportion of all predictions that were correctly classified.
- **Sensitivity:** the proportion of true positive cases (ERCC2-MUT) that were correctly identified by the model. High sensitivity indicates effective detection of mutation-positive cases.
- **Specificity:**: the proportion of true negative cases (ERCC2-WT) that were correctly identified. High specificity reflects the ability to avoid false positives.
- **Positive Predictive Value (Precision)**: the proportion of predicted positives that were true positives.
- **Negative Predictive Value:** the proportion of predicted negatives that were true negatives.
- **Balanced Accuracy:**the average of sensitivity and specificity, which is useful in cases of class imbalance.

### 2 Supplemental Figures

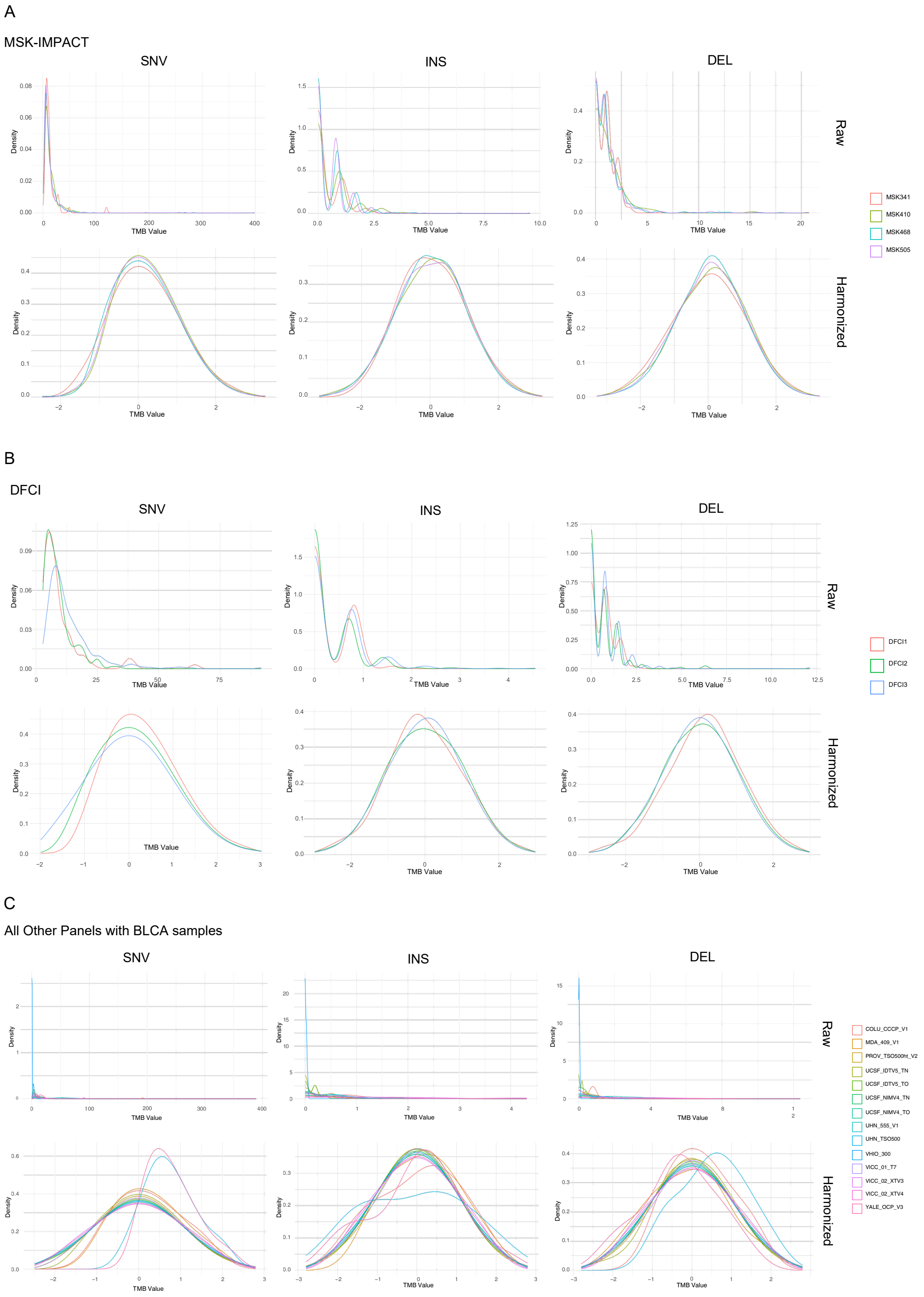


**Supplemental Figure 1:** Distribution of raw (top) and harmonized (bottom) TMB values for SNV, INS and DEL in **(A)** MSK-IMPACT **(B)** DFCI-ONCOPANEL and **(C)** All other panels with BLCA samples.

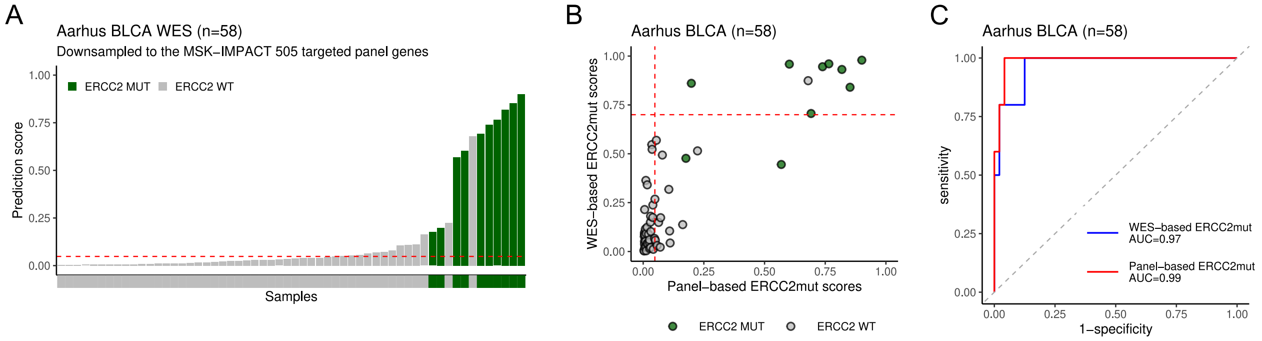


**Supplemental Figure 2:** **A.** Distribution of panel-based ERCC2mut scores in the Aarhus WES cohort calculated using mutations downsampled to genes present in the MSK-IMPACT 505 panel. **B.** Correlation between the panel-based and WES-based ERCC2mut scores in the Aarhus cohort. The red dashed lines indicate the cut-off values used to separate ERCC2-MUT and WT cases: 0.048 for the panel-based model (x axis) and 0.7 for the WES-based model (y axis). **C.** ROC curve and AUC for the panel-based and WES-based ERCC2mut models in the Aarhus cohort.

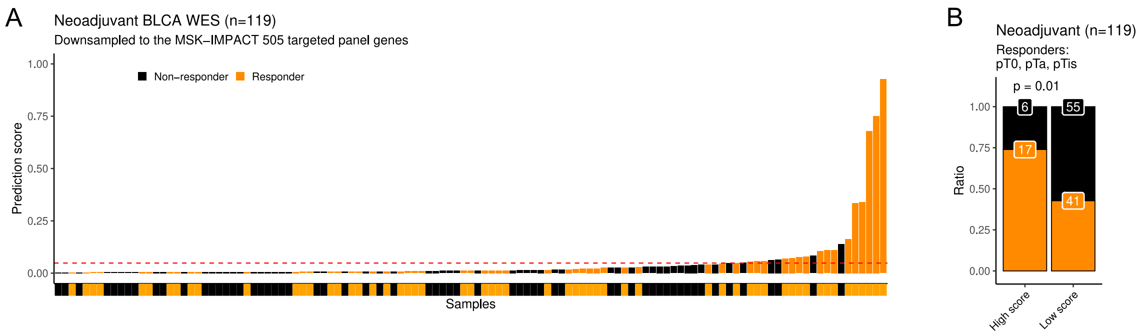


**Supplemental Figure 3:** **A.** Distribution of panel-based ERCC2mut scores in the combined neoadjuvant muscle-invasive bladder cancer (MIBC) cohort excluding pathogenic ERCC2-MUT cases (only ERCC2-WT cases) colored by response to neoadjuvant platinum-based chemotherapy. **B.** Bar plots showing the ratio of responders and nonresponders among ERCC2-WT cases with high and low panel-based ERCC2mut score (cut-off value = 0.048)s in the neoadjuvant MIBC cohort excluding pathogenic ERCC2-MUT cases (Fisher's exact test: p-value = 0.01). Response was defined as pT0, pTa, pTis, and pT1.

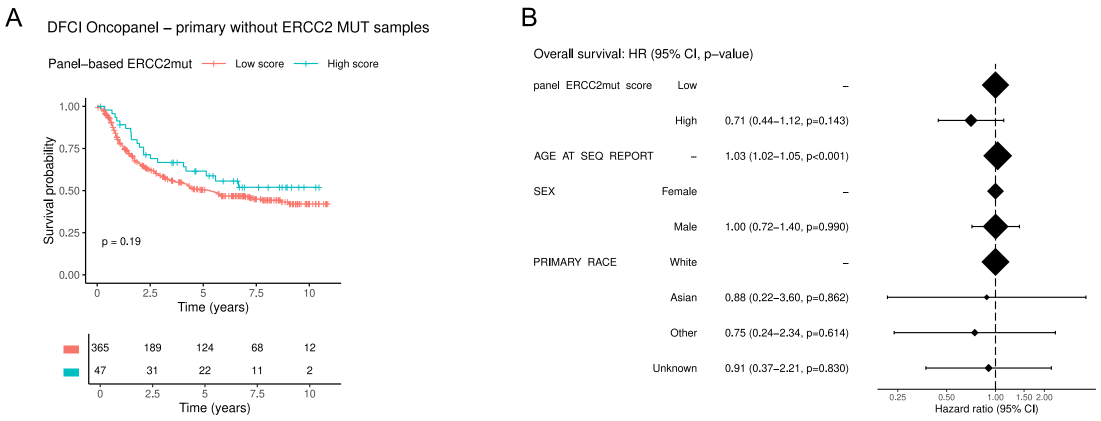


**Supplemental Figure 4:** A. Kaplan-Meier survival curves and risk table for BLCA cases with primary samples sequenced in the DFCI-Oncopanel dataset, excluding known pathogenic *ERCC2* mutations, stratified by the panel-based ERCC2mut prediction score (cut-off: 0.128; log-rank test: p-value = 0.19). **B.** Hazard ratios with 95% CIs and p-values estimated using the Cox proportional hazards model (CPH) model.

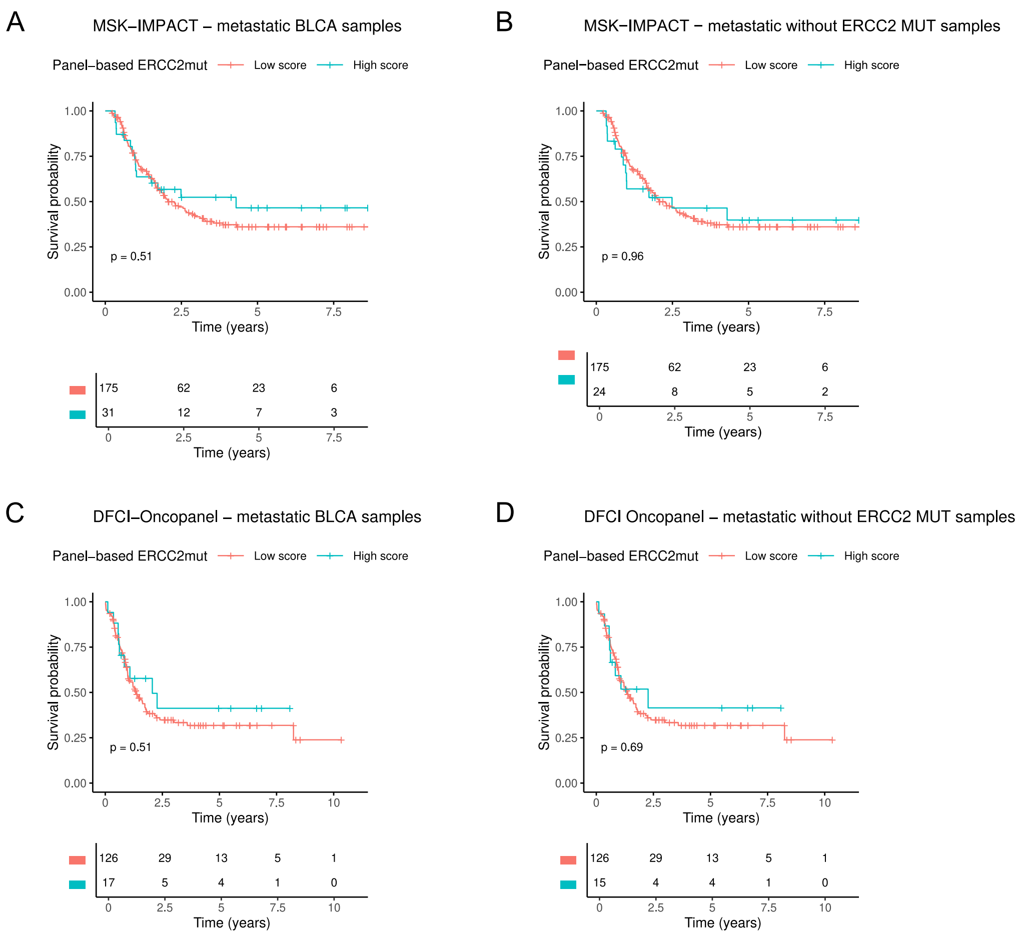


**Supplemental Figure 5:** Kaplan-Maier survival curves of patients with samples from metastatic sites in the MSK-IMPACT (A-B) and DFCI-Oncopanel (C-D) BLCA subsets of the GENIE cohort.

### 3 Supplemental Tables

| ERCC2-MUT |
|---|
| Y14C |
| Y24C |
| M42V |
| S44L |
| T46S |
| Y72C |
| E79D |
| E86Q |
| Y209C |
| N238S |
| V242F |
| S246F |
| S246Y |
| P463L |
| T484M |
| E576K |
| E606G |
| G607A |
| D609E |
| D609G |
| H659Y |
| G665A |
| G665C |
| E606Q |

**Supplemental Table 1:** List of experimentally tested cisplatine-sensitive *ERCC2* mutations [5, 6] used to determine *ERCC2* mutation status of each sample.

| Cohort | Dataset | Type | ERCC2-MUT | ERCC2-WT | Total |
|---|---|---|---|---|---|
| GENIE | MSK-IMPACT | Training | 114 | 1495 | 1787 |
| GENIE | DFCI Oncopanel | Test | 26 | 523 | 623 |
| | Pooled panels with BLCA samples | Test | 22 | 476 | 548 |
| | Pathogenic *ERCC2* variants in other cancer types | Test | 44 | 0 | 44 |
| DFCI-MSKCC | DFCI-MSKCC Neoadjuvant | Test | 9 | 41 | 50 |
| Aarhus | Aarhus Neoadjuvant | Test | 12 | 48 | 60 |
| Philadelphia | Philadelphia Neoadjuvant | Test | 10 | 38 | 48 |

**Supplemental Table 2:** Number of ERCC2-MUT, ERCC2-WT, and total samples in the training (MSK-IMPACT) and validation datasets.

| Panel | ERCC2-MUT | ERCC2-WT | Total |
|---|---|---|---|
| COLU-CCCP-V1 | 0 | 6 | 6 |
| MDA-409-V1 | 0 | 10 | 11 |
| PROV-TSO500HT-V2 | 13 | 153 | 189 |
| UCSF-IDTV5-TN | 2 | 66 | 73 |
| UCSF-IDTV5-TO | 4 | 67 | 76 |
| UCSF-NIMV4-TN | 0 | 42 | 43 |
| UCSF-NIMV4-TO | 0 | 15 | 17 |
| UHN-TSO500-V1 | 0 | 9 | 10 |
| UHN-555-V1 | 3 | 15 | 19 |
| VICC-01-T7 | 0 | 40 | 43 |
| VICC-02-XTV3 | 0 | 6 | 7 |
| VICC-02-XTV4 | 0 | 9 | 10 |
| VHIO-300 | 0 | 24 | 29 |
| YALE-OCP-V3 | 0 | 14 | 15 |
| Total | 22 | 476 | 548 |

**Supplemental Table 3:** Pooled targeted panels from the GENIE cohort with BLCA samples and their respective number of ERCC2-MUT, ERCC2-WT, and total samples sequenced with the respective panel. Only panels with $> 5$ BLCA samples and where each sample has $\geq 5$ mutations were included.

| Dataset | Optimal cutpoint |
|---|---|
| MSK-IMPACT | 0.04821027 |
| DFCI-ONCOPANEL | 0.1279685 |
| Pooled panels with BLCA samples | 0.1322924 |

**Supplemental Table 4:** Optimal cutpoint values for the training and validation datasets of the panel-based ERCC2mut classifier.

| Targeted panel version | Number of genes |
|---|---|
| MSK-IMPACT 341 | 341 |
| MSK-IMPACT 410 | 410 |
| MSK-IMPACT 468 | 468 |
| MSK-IMPACT 505 | 505 |
| DFCI Oncopanel 1 | 304 |
| DFCI Oncopanel 2 | 326 |
| DFCI Oncopanel 3 | 447 |

**Supplemental Table 5:** Number of genes included in the MSK-IMPACT and DFCI Oncopanel targeted sequencing panels.

| | **Reference** | |
|---|---|---|
| **Prediction** | 0 (ERCC2-WT) | 1 (ERCC2-MUT) |
| 0 (ERCC2-WT) | 1327 | 2 |
| 1 (ERCC2-MUT) | 168 | 112 |

**Supplemental Table 6:** Confusion matrix for the performance of the panel-based ERCC2mut classifier on the MSK-IMPACT training set.

| | **Reference** | |
|---|---|---|
| **Prediction** | 0 (ERCC2-WT) | 1 (ERCC2-MUT) |
| 0 (ERCC2-WT) | 481 | 4 |
| 1 (ERCC2-MUT) | 42 | 22 |

**Supplemental Table 7:** Confusion matrix for the performance of the panel-based ERCC2mut classifier on the DFCI-ONCOPANEL testing set.

| | **Reference** | |
|---|---|---|
| **Prediction** | 0 (ERCC2-WT) | 1 (ERCC2-MUT) |
| 0 (ERCC2-WT) | 423 | 4 |
| 1 (ERCC2-MUT) | 53 | 18 |

**Supplemental Table 8:** Confusion matrix for the performance of the panel-based ERCC2mut classifier on the testing set made from all panels, beyond the MSK-IMPACT and DFCI-Oncopanel panels, within the GENIE cohort with BLCA samples.

**ERCC2mut**

| Model | WES-based [7] | Panel-based |
|---|---|---|
| Algorithm | logistic regression | gradient boosting classifier combining decision trees |
| Predictor features | ID8 | Cosine similarity with ID8 |
| | SBS5 (Signature 5) | Cosine similarity with SBS5 |
| | SBS2 (Signature 2) | Cosine similarity with SBS2 |
| | - | Cosine similarity with SBS13 |
| | - | SNV TMB |
| | - | deletion TMB |
| | ID10 | - |
| | ID2 | - |
| | DBS4 | - |
| | TSB ratio T>A | - |
| | TSB ratio C>G | - |
| | TSB ratio T>G | - |
| Cut-off value (training dataset) | 0.7 | 0.04821027 |

**Supplemental Table 9:** Details regarding the WES-based and panel-based ERCC2mut models.

| DFCI-MSKCC | ERCC2-MUT | | ERCC2-WT | |
|---|---|---|---|---|
| panel-based / WES-based | 0 | 1 | 0 | 1 |
| 0 | 1 | 0 | 30 | 1 |
| 1 | 0 | 8 | 5 | 4 |

**Supplemental Table 10:** Comparison of the predictions of the panel- and WES-based ERCC2mut models in the down-sampled DFCI-MSKCC BLCA WES cohort.

| Philadelphia | ERCC2-MUT | | ERCC2-WT | |
|---|---|---|---|---|
| panel-based / WES-based | 0 | 1 | 0 | 1 |
| 0 | 3 | 2 | 34 | 3 |
| 1 | 0 | 5 | 0 | 1 |

**Supplemental Table 11:** Comparison of the predictions of the panel- and WES-based ERCC2mut models in the down-sampled Philadelphia BLCA WES cohort.

| Aarhus | ERCC2-MUT | | ERCC2-WT | |
|---|---|---|---|---|
| panel-based / WES-based | 0 | 1 | 0 | 1 |
| 0 | 0 | 2 | 33 | 14 |
| 1 | 0 | 8 | 0 | 1 |

**Supplemental Table 12:** Comparison of the predictions of the panel- and WES-based ERCC2mut models in the down-sampled Aarhus BLCA WES cohort neoadjuvant subset.

| NAC Cohorts | Sequencing | Total number of samples | Total number of samples after downsampling | ERCC2-MUT |
|---|---|---|---|---|
| DFCI-MSKCC | WES | 50 | 49 | 9 |
| Philadelphia | WES | 48 | 48 | 10 |
| Aarhus | WES | 60 | 58 | 10 |
| GENIE | MSK-IMPACT | 38 | Not applicable | 37 |
| Total | | 196 | 193 | 66 |

**Supplemental Table 13:** Combined neoadjuvant cohort.

[3] Max Kuhn. *caret R package*. https://topepo.github.io/caret/.

[4] Greg Ridgeway. *gbm3 R package*. https://github.com/gbm-developers/gbm3.

[5] Qiang Li et al. "ERCC2 helicase domain mutations confer nucleotide excision repair deficiency and drive cisplatin sensitivity in muscle-invasive bladder cancer". In: *Clinical Cancer Research* 25 (3 2019), pp. 977–988. ISSN: 15573265. DOI: 10.1158/1078-0432.CCR-18-1001.

[6] Judit Börcsök et al. "Quantitative functional profiling of ERCC2 mutations deciphers cisplatin sensitivity in bladder cancer". In: *Journal of Clinical Investigation* 135 (16 Aug. 2025). ISSN: 1558-8238. DOI: 10.1172/JCI186688.

[7] Judit Börcsök et al. "Identification of a synthetic lethal relationship between nucleotide excision repair deficiency and irofulven sensitivity in urothelial cancer". In: *Clinical Cancer Research* 27 (7 Nov. 2021), pp. 2011–2022. ISSN: 15573265. DOI: 10.1158/1078-0432.CCR-20-3316. URL: http://www.ncbi.nlm.nih.gov/pubmed/33208343.
